# Mmp10 is required for post-translational methylation of arginine at the active site of methyl-coenzyme M reductase

**DOI:** 10.1101/211441

**Authors:** Zhe Lyu, Chau-wen Chou, Hao Shi, Ricky Patel, Evert C. Duin, William B. Whitman

## Abstract

Catalyzing the key step for anaerobic methane production and oxidation, methyl-coenzyme M reductase or Mcr plays a key role in the global methane cycle. The McrA subunit possesses up to five post-translational modifications (PTM) at its active site. Bioinformatic analyses had previously suggested that methanogenesis marker protein 10 (Mmp10) could play an important role in methanogenesis. To examine its role, MMP1554, the gene encoding Mmp10 in *Methanococcus maripaludis*, was deleted with a new genetic tool, resulting in the specific loss of the 5-(S)-methylarginine PTM of residue 275 in the McrA subunit and a 40~60 % reduction in the maximal rates of methane formation by whole cells. Methylation was restored by complementations with the wild-type gene. However, the rates of methane formation of the complemented strains were not always restored to the wild type level. This study demonstrates the importance of Mmp10 and the methyl-Arg PTM on Mcr activity.

## Introduction

Methyl-coenzyme M reductase or Mcr catalyzes the last step of methane formation in all methanogenic Archaea and the first step of anaerobic methane oxidation in the methanotrophic Archaea or ANME (1, 2). It catalyzes the reversible reaction shown below that results in production of 500-600 Tg of CH_4_ and oxidation of 70-300 Tg of CH_4_ per year (3).

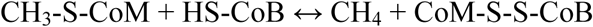

where CH_3_-S-CoM is methyl-coenzyme M and HS-CoB is coenzyme B. CH_4_ is an important biofuel as well as a potential feedstock for the chemical industry if it could be converted by Mcr to a liquid biofuel with a high energy density (4, 5). CH_4_ is also a potent greenhouse gas, increases of which are contributing to global warming (3). Therefore, understanding the biochemistry of Mcr is significant not only to advance a CH_4_-based bioeconomy but also to develop environmental CH_4_ mitigation strategies.

Although Mcr is notoriously difficult to study due to its extreme sensitivity to oxygen and lability of the active reduced form, structural and biochemical studies have uncovered many unique aspects (1, 6, 7). Using the *Methanothermobacter marburgensis* Mcr as a model, it has been shown that Mcr is a hexameric, 300 kD protein composed of three different subunits in an α_2_β_2_γ_2_ configuration (1). It contains two molecules of an unusual Ni tetrapyrrole, coenzyme F_430_ or methylthio-F_430_ in ANME-1, which is tightly but not covalently bound (1, 6). During enzymatic catalysis, the Ni(I) attacks the sulfur atom of methyl coenzyme M, producing a methyl radical intermediate, as proposed recently (7). The enzyme possesses two identical active sites, each of which contains up to five post-translationally modified amino acid residues(1). In *M. maripaludis* (Z. Lyu, CW. Chou, H. Shi, L. Wang, R. Ghebreab, D. Philllips, Y. Yan, E. C. Duin and W. B. Whitman, submitted for publication), the post-translational modifications (PTMs) include (*M. maripaludis* numbering) 1-*N*-methyl-His^261^, 5-(*S*)-methyl-Arg^275^, 2-(*S*)-methyl-Gln^403^, and thio-Gly^448^, which are all found within the McrA subunit (1). The S-methylation of cysteine, which is common in the Mcr of many methanogens (8), is in low abundance or absent in *M. maripaludis*. Generally, the PTMs are highly conserved among the Mcr from methanogens if not the ANME (8). For instance, the arginine methylation has been found in all methanogenic Mcr examined but not the ANME-1 Mcr.

Given the wide occurrence but limited diversity of these PTMs across the Mcr from distantly related species, they are believed to be important to catalysis. However, little is known about their biosynthesis or function. S-adenosyl methionine is believed to be the methyl donor for the methylations (9). Very recently, *tfuA* and *ycaO* (or methanogen marker protein 1) homologs were found to be required for the thio-Gly PTM in *Methanosarcina acetivorans* (10). The present study identified a gene required for the 5-(*S*)-methyl-Arg PTM in *M. maripaludis*, a genetically tractable model organism for methanogenic Archaea (11).

## Results

### Mmp10 is a candidate Arg methyltrasnferase for PTMs

Methanogens encode a large number of hypothetical methyltransferases with unknown specificity, making it difficult to identify candidates for the PTMs of Mcr (9, 12). However, several criteria may help with identification of potential candidates. First, candidates for 5-(*S*)-methyl-Arg and N-methyl-His methyltransferase genes were hypothesized to be widely distributed among methanogens as these two modifications are conserved in all methanogens examined so far. Second, the methyltransferases were expected to be absent in most if not all non-methanogenic Archaea, which do not encode a Mcr. Third, the 5-(*S*)-methyl-Arg methyltransferase was expected to be absent in the ANME-1 methanotrophs, which lack this PTM. Fourth, this methyltransferase should possess a SAM-binding site, which is essential for SAM-dependent methylation of Mcr. Lastly, the methyltransferase gene might be located close to the *mcr* operon so that expression, modification and folding of Mcr would be better coordinated.

Classified into the TIGR03278 family by ProPhylo, the methanogenesis marker protein 10 (Mmp10) seemed to meet all of the aforementioned criteria. It was found in many methanogens in a previous bioinformatic study and likely encodes a radical SAM enzyme, which has been long assumed to play an important role in methanogenesis (13). Located next to but transcribed divergently from the *mcr* operon (**Fig. 1A**), MMP1554 is the *mmp10* homolog in *M. maripaludis* (14). Bioinformatic analysis here with a much larger dataset further confirmed that *mmp10* homologs were widely distributed in all methanogen lineages except *Methanoculleus bourgensis* and *Methanomassiliicoccales* spp. (**Table 1**). Homologs were also found in ANME-2 methanotrophs and two unclassified euryarchaeotes but not in ANME-1 and any of the other archaeal species examined. These included the *Candidatus* ‘Bathyarchaeota’ and ‘Verstraetearchaeota’ genomes where other methanogenesis genes were recently found (15, 16). Among methanogens and methanotrophs that possess *mmp10* homologs, it was adjacent to the *mcr* operon except in the orders *Methanomicrobiales* and *Methanocellales*. Therefore, it was concluded that Mmp10 would be a reasonable candidate for a Mcr methyltransferase.

**Fig. 1.**
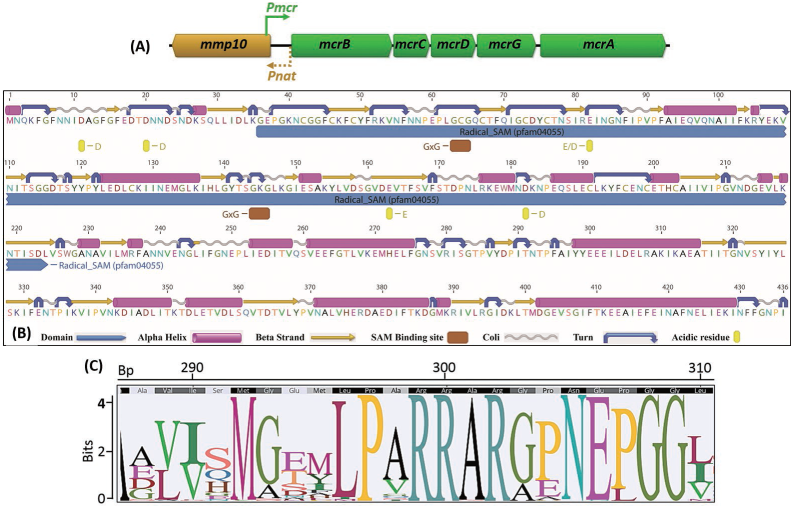
Structural features of Mmp10 and its potential target sequence in McrA. (**A**) In *M. maripaludis*, Mmp10 is encoded by MMP1554, which is transcribed divergently from the adjacent *mcrBCDGA* operon that encodes the methyl-coenzyme M reductase (14). The intergenic sequence between *mmp10* and *mcr* is only 259 bp in length. Therefore, *Pmcr*, the ~290 bp long promoter that drives *mcr* expression (11), extends partially into the coding region of *mmp10*. The exact promoter sequence for *mmp10* is unknown, but B recognition element (BRE) and TATA box sequences could be predicted from within the intergenic sequence. Thus, the whole intergenic sequence is taken as the predicted *mmp10* native promoter namely *Pnat* in this study. (**B**) Structural features predicted for the Mmp10 protein. SAM-dependent methyltransferases have a highly conserved GxG motif in the first β-sheet and an acidic residue (D or E) in the second β-sheet. While β-sheets are difficult to predict without a crystal structure, β-strands and turns that form β-sheets can be more easily identified. Two GxG motifs (brown boxes) were found. The one positioned at 144~146 was in a region likely to form a β-sheet and conserved in all homologs from six genomes representing each methanogen order and one ANME-2 genome (**Fig. S1**). Both upstream and downstream of the conserved GxG motif, multiple conserved acidic residues (yellow boxes) could be also identified. At least three of them could be located in a β-sheet, at positions 20, 82 and 163. Both positions 82 and 163 were conserved among the homologs, except that the position 163 of *Methanopyrus kandleri* was a nucleophilic S residue instead of the conserved acidic E residue (**Fig. S1**). The legend for each structural feature is shown at the bottom of the figure. (**C**) Sequence logo of the context of Met-Arg from an alignment of 251 McrA amino acid sequences from all seven orders of methanogens available on the IMG at the time of analysis. On top of the logo, sequence consensus and coordinates are shown, and Met-Arg is located at position 300 in this alignment.

**Table 1.** Distribution of the methanogenesis marker protein 10 (Mmp10) homologs within the Archaea

| Lineages | No. of genomes | No. of genomes encoding Mmp10 | No. of genomes with *mmp10* flanking *mcr*<sup>a</sup> |
| --- | --- | --- | --- |
| *Methanococcales* | 19 | 19 | 19 |
| *Methanopyrales* | 1 | 1 | 1 |
| *Methanobacteriales* | 58 | 58 | 53<sup>b</sup> |
| *Methanomicrobiales* | 20 | 19<sup>c</sup> | 0 |
| *Methanocellales* | 3 | 3 | 0 |
| *Methanosarcinales* | 112 | 109<sup>d</sup> | 98<sup>e</sup> |
| *Methanomassiliicoccales* | 8 | 0 | 0 |
| Non-methanogenic *Euryarchaeota* | 275 | 0 | 0 |
| Unclassified *Euryarchaeota* | 75 | 2<sup>f</sup> | 0 |
| *Bathyarchaeota* | 24 | 0 | 0 |
| *Verstraetearchaeota* | 5 | 0 | 0 |
| Other Archaea | 359 | 0 | 0 |

<sup>a</sup>Marker 10 position was examined by "Show neighborhood regions with the same top COG hit (via top homolog)" command at IMG/M ER.
<sup>b</sup>Not linked to *mcr* in *Methanobrevibacter smithii* TS145B and ACE6, *Methanobrevibacter curvatus* DSM11111, and *Methanobacterium* sp. Maddingley. These are all draft genomes where *mmp10* and *mcr* are on different contigs; in *Methanosphaera stadtmanae*, *mmp10* flanks the *mcrC*-*mtr* operon with *mrtBDGA* 16.8kb downstream of the *mcr* operon.
<sup>c</sup>Absent in *Methanoculleus bourgensis* BA1.
<sup>d</sup>Absent in *Methanosarcina horonobensis* JCM 15518 (draft genome), *Methanosarcina mazei* Tuc01; and ANME-1 (FP565147); present in all five ANME-2 genomes.
<sup>e</sup>Not linked in four of the five ANME-2 genomes, all three *Methanosaeta* genomes, *Methanosarcina mazei* 1.H.A.2.8 (draft genome with *mmp10* and *mcr* on separate contigs), *Methanosalsum zhilinae*, *Methanoflorens stordalenmirensis*, and *Methermicoccus shengliensis*.
<sup>f</sup>Present in *Euryarchaeota* archaeon JGI 0000059-G05 and *Euryarchaeota* sp. AmaM. Both *rRNA* and *mcrA*, the conventional gene markers, were absent in the first archaeon. However, it encoded an McrC subunit which had 85% of amino acid identity to that from *Methanosaeta concilli*. The second archaeon has been recently named *Methermicoccus shengliensis* strain AmaM (51). Therefore, both archaeal species are in fact methanogens.

This conclusion was supported by an examination of the predicted domain and coenzyme binding sites for Mmp10 (**Fig. 1B**). Mmp10 has a SAM domain (pfam04055) of 190 amino acids on its N-terminus. Although its function is unknown, the similarly sized C-terminus is also highly conserved. This agrees with the structure of known methyltransferases, which typically consist of well-conserved SAM domains but different substrate-binding domains (17). This structural feature was conserved in nearly all Mmp10 homologs examined. A few exceptions came from draft genomes including *Methanobacterium formicicum* JCM 10132 (Ga0128400_10744), *Methanobrevibacter arboriphilus* JCM 13429 (Ga0128401_11159), *Methanobrevibacter oralis* JMR01 (Ga0053424_102302), *Methanogenium cariaci* JCM 10550 (Ga0128318_10529), *Methanosarcina barkeri* JCM 10043 (Ga0128387_101358), *Methanosarcina mazei* JCM 9314 (Ga0128314_100641) and an ANME-2 (ANME2D_draft_0001.00000480). For these genes, either the N- or C-termini were truncated, possibly because the genomes were not complete. Indeed, closely related genomes of the same genera or species all possessed the complete *mmp10*. The exception was *M. cariaci*, which was the only *Methanogenium* genome available in the IMG database.

Although SAM-dependent methyltransferases are very diverse, they generally share a highly conserved GxG motif that binds to the SAM nucleotide and an acidic residue located downstream that forms hydrogen bonds with the hydroxyl groups of the SAM ribose (18). The GxG motif is found at the end of one β-sheet, whereas the acidic residue is located in another β-sheet (18). Two candidates for this motif and downstream acidic residue were also present in the Mmp10 (**Fig. 1B and Fig. S1 in supplementary material**).

To gain insight into a possible recognition sequence for the methyltransferase, the R^α275^ region of the methanogen and ANME genomes available in the IMG database were compared by Sequence Logo (**Fig. 1C**). Because the McrA subunit is highly conserved near the active site, the region near R^275^ contained a large number of invariant residues. However, limiting a possible methylation consensus sequence to three residues on either side of the methylation site yielded a candidate motif of PxR^274^R^275^(A/S)R(G/A). Within this motif, the unmodified R^274^ could interact with the coenzyme B at the active site due to their close interatomic distances, as revealed by a recent published methanococcal MCR crystal structure (**Fig. 2**) (19). While the methylated R^275^ is located far away from coenzyme B, the methylation may provide structural benefits that could stabilize and/or enhance interactions between the R^274^ and coenzyme B.

**Fig. 2.**
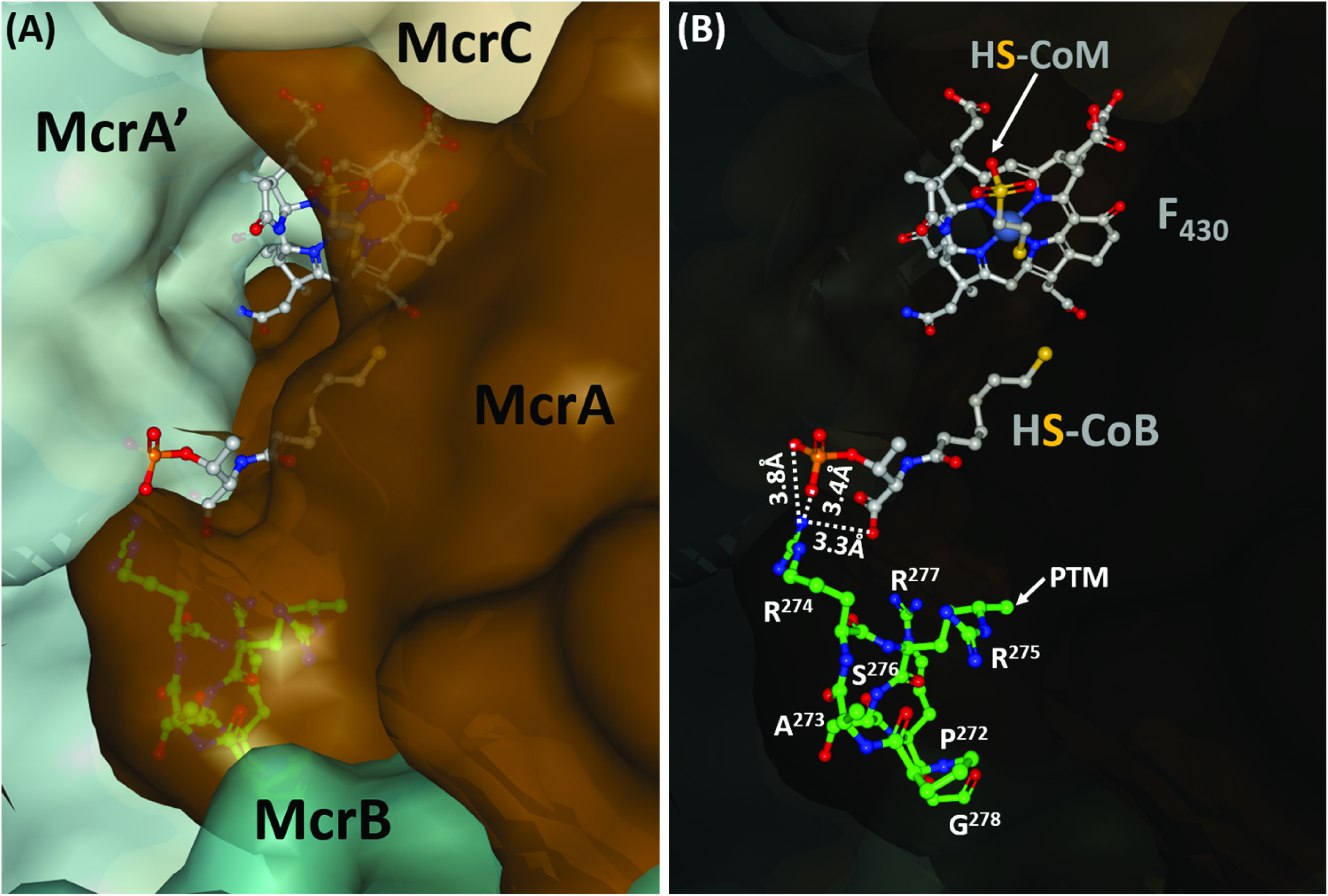
Methanococcal Mcr active site as illustrated by the crystal structure of *Methanothermococcus thermolithotrophicus* (5N1Q) (19). **(A)** View from the surface of Mcr. Shown in gray ball-and-stick model, the coenzymes F_430_, M (HS-CoM), and B (HS-CoB) are embedded within the active site pocket created primarily by the McrA and McrA’ subunits. **(B)** A see-through view shows the PAR^274^R^275^SRG motif (shown in green ball-and-stick model) of McrA contributing to the formation of the active site pocket. The post-translational modified (PTM) methyl group is indicated for R^275^, which is located far away from the HS-CoB, e.g., the distance between the carbon atom of this PTM and the negatively charged oxygen atoms of the carboxyl group on the HS-CoB is 6.7 Å. In contrast, its neighboring R^274^ is much closer to the HS-CoB, suggesting potential interactions. The positively charged nitrogen atom of the amine group of R^274^ and the negatively charged oxygen atoms of the phosphate or carboxyl groups of the HS-CoB were within 3.3-3.8 Å of each other, as indicated by the dash lines.

### A new tool for markerless deletion of mmp10

A new plasmid vector p5L-R was constructed by assembling standardized genetic modules through BioBrick assembly (**Fig. S2**). The standard modules included a pUC57 backbone, methanococcal promoter, selectable markers, methanococcal ribosomal binding sites (RBS), and repetitive elements (RE), which were all BioBrick compatible. The pUC57 backbone harbored the origin of replication and *amp* for selection in *E. coli*. The positive and negative selectable markers in *Methanococcus* were *pac* and *hpt*, genes conferring puromycin resistance and 8-azahypoxanthine sensitivity, respectively (20, 21). They were synthesized to optimize codon usage for *pac*, remove extraneous bases, and eliminate internal BioBrick sites for *hpt*. PCR amplified from the pAW42 vector (22), the strong and constitutive promoter *PhmvA* drove expression of the selectable markers. To minimize homologous recombination between identical RBSs, different RBSs were introduced at the 5’ ends of *pac* and *hpt* through PCR. They were the 14 bp sequences immediately upstream of the *mcrB* and *hmvA* genes from *Methanococcus voltae*, respectively. In addition, a short sequence downstream of the *Methanococcus voltae mcr* operon which contained a terminator (Tmcr) was also introduced at the 3’-end of the *hpt-pac* cassette. The selectable markers were then flanked with two identical REs forming a RMR module, i.e., RE-Markers-RE, of which the REs can undergo homologous recombination to remove the markers and leave a short scar (**Fig. S3**). This marker removal strategy has been described recently, suggesting RE size affects homologous recombination efficiency (23). The p5L-R also included the upstream and downstream sequences of MMP0148, which were used to test its effectiveness for construction of the markerless deletion of MMP0148. Of the various RE sizes of 20, 40, 60, 120 and 180 bp examined, a minimal RE of 40 bp was needed for detectable marker removal. With a RE of 60 bp, marker deletion was always observed when making several mutants, including the *∆mmp10* deletion (see below) in *M. maripaludis*.

The full protocol, standardized primers and plasmid sequence for markerless deletions using p5L-R can be found in **Supplementary Methods**. Briefly, The RMR module of the p5L-R was PCR amplified with standardized primers with a different SfiI overhang at each end (**Table S1**). Then 0.5~1 kb of upstream and downstream sequences of the target gene were PCR amplified with the same overhangs at the 3’-end of the upstream and 5’-end of the downstream sequence, respectively. The PCR products were digested with SfiI and ligated to create the initial construct, which was either used directly for transformation of *M. maripaludis* or PCR amplified to increase the amount of DNA. The p5L-R procedure (**Fig. S3**) addressed several shortcomings of previous markerless deletion tools in *M. maripaludis* (24). One, this strategy allowed for very reliable removal of the wild type genotype, while the previous tool yielded large numbers of wild type cells which had to be removed by screening (24). Two, the *M. maripaludis* genome does not contain any SfiI sites, so this strategy can be used for deletion of any gene. In contrast, the previous tools possessed multiple cloning sites that overlapped with restriction sites in the genome (24). Third, low concentrations of antibiotic (1 to 2.5 μg mL^−1^ of puromycin) were effective for positive selection, while high concentrations of neomycin (500 to 1000 μg mL^−1^) were necessary with the previous tools (24). Lastly, the non-palindromic restriction enzyme SfiI minimized self-ligation, and cloning in *E. coli* was not required (25). Previously, two rounds of cloning were needed to make the final construct (24).

### Mmp10 was necessary for arginine PTM

*M. maripaludis* mutants were constructed to investigate the role of Mmp10 in PTMs of Mcr. Strain S0030 possessed a *∆mmp10* deletion in the S0001 background. In strain S0031, the *∆mmp10* mutation was complemented with the *mmp10* gene expressed under the control of *Pnat*, the predicted native promoter, from the plasmid pM10. In strain S0034, the deletion was complemented with *mmp10* expressed under the control of *PhmvA*, a constitutively strong promoter commonly used for *M. maripaludis* (26–28).

Mcr proteins were purified from the wild-type and mutant strains to determine the PTMs by LC MS/MS analysis. Three of the PTMs, 1-*N*-methyl-His^261^, 2-(*S*)-methyl-Gln^403^, and thio-Gly^448^ of McrA, were found in the enzyme from the wild type and *∆mmp10* strains (data not shown). In partial trypsin digestions of the Mcr from wild-type S0001 cells, the 5-(*S*)-methyl-Arg modification was observed within an Arg-rich region of the protein in the peptide H*ADVIQMGNALPGRr^275^, where H* is the methyl-His^261^ in a position homologous to that found in other methanogens and r^275^ is the methyl-Arg. Because the methylation was on the second Arg in the peptide, it was only observed in partial trypsin digestions. In the McrA from S0001, the partially digested peptides containing methyl- r^275^ represented only a small fraction of the peptides covering this region, about 3 % of the abundance of the fully digested peptide (**Table S2 and Fig. S4**). In contrast, the peptide containing the methyl-Arg PTM was not observed in the enzyme from the deletion strain S0030, even though the unmodified RR form of the partial digestion product increased to 8 % of the fully digested peptide (**Table S2 and Fig. S5**). Tryptic digestions tend to be incomplete at dibasic sites such as RR, RK and KK (29, 30). Thus, removal of the Arg modification was expected to increase the rate of partial cleavage as observed. In the Mcr from the complementation strain S0031, the unmodified peptide was not observed, and the fraction of partially digested peptide was reduced again to 4 %, as seen in the wild-type (**Table S2 and Fig. S6**).

While these results suggested that *mmp10* was required for the PTM of Arg^275^, the partial digestion by trypsin made quantification of the extent of modification difficult. For that reason, a pepsin degradation was developed that yielded the peptide PGRr^275^ARGPNEPGGIRF and enabled unambiguous quantification of the methyl-Arg^275^. Based on peak area integration, Me-Arg^275^ was absent in the Mcr of the *∆mmp10* deletion strain (**Table 2 and Fig. S7**). In contrast, more than 96-98% of the Arg^275^ was methylated in the complementation and wild type strains (**Table 2 and Fig. S8 to S10**). These results confirmed that *mmp10* was required for the Arg^275^ methylation.

**Table 2.**
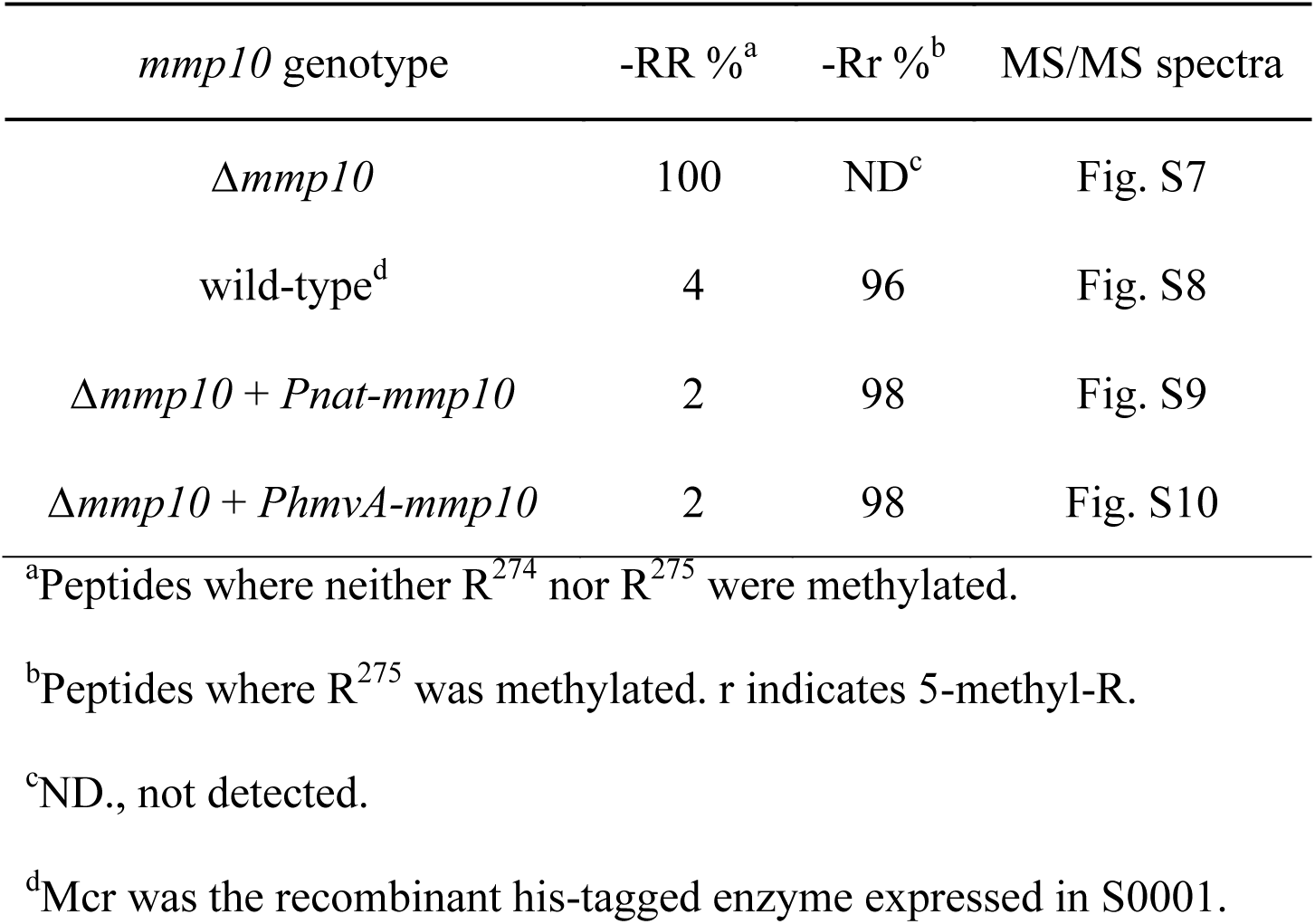
Abundance of peptides containing methyl-R^α275^ after pepsin digestion

| *mmp10* genotype | -RR %<sup>a</sup> | -Rr %<sup>b</sup> | MS/MS spectra |
| --- | --- | --- | --- |
| *Δmmp10* | 100 | ND<sup>c</sup> | Fig. S7 |
| wild-type<sup>d</sup> | 4 | 96 | Fig. S8 |
| *Δmmp10 + Pnat-mmp10* | 2 | 98 | Fig. S9 |
| *Δmmp10 + PhmvA-mmp10* | 2 | 98 | Fig. S10 |

<sup>a</sup>Peptides where neither R<sup>274</sup> nor R<sup>275</sup> were methylated.
<sup>b</sup>Peptides where R<sup>275</sup> was methylated. r indicates 5-methyl-R.
<sup>c</sup>ND., not detected.
<sup>d</sup>Mcr was the recombinant his-tagged enzyme expressed in S0001.

### Growth phenotype of ∆mmp10 mutants

The growth of the *∆mmp10* deletion strain S0030 was severely inhibited compared to the wild type. Not only did the lag phase increase, but the growth rate was reduced from 0.27 + 0.03 h^−1^ (average + standard deviation, n = 3) in the wild-type to 0.17 ± 0.03 h^−1^ in S0030 (**Fig. 3**). Complementation of the deletion failed to restore wild-type growth. Strain S0031 had a prolonged lag and its growth rate was further reduced to 0.14 + 0.01 h^−1^. While strain S0034 grew somewhat better than the deletion mutant, the lag was longer and the growth rate, 0.19 ± 0.03 h^−1^, was still slower than wild type. To examine if the levels of Mcr expression were different among the strains, whole cell extracts were separated by SDS PAGE (**Fig. 4**). Although the McrA subunit was not well separated from other bands, the McrB and McrG subunits were readily identifiable, and the Mcr abundance for each strain could be calculated based on integration of the band intensity. Similar levels of Mcr were found in triplicates of all strains (*p* = 0.13 by One-way ANOVA), and they were 13.6 ± 1.8 % (average and standard deviation of three measurements, S0001, the wild type), 14.2 ± 1.2 % (S0030, the Δ mutant), 14.5 ± 1.0 % (S0034, the *PhmvA*-*mmp10* complement) and 12.6 ± 1.5 % (S0031, the *Pnat*-*mmp10* complement). The similar SDS profiles among the strains also suggested similar expression levels for most other proteins (**Fig. 4**). Thus, the *∆mmp10* deletion and complementation did not appear to affect the levels of Mcr.

**Fig. 3.**
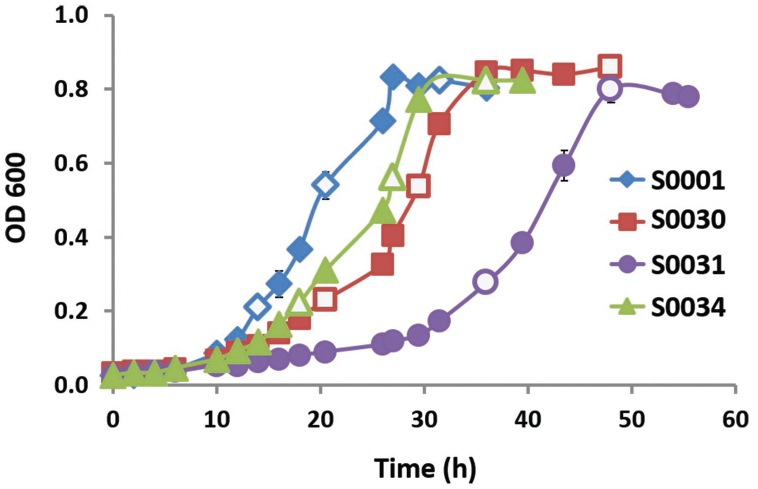
Growth of the *∆mmp10* mutant and complementation strains. *M. maripaludis* strains S0001 (the wild-type), S0030 (the deletion mutant), S0031 (the *Pnat-∆mmp10* complement), and S0034 (the *PhmvA-∆mmp10* complement). Cultures were grown in 20 mL of complex formate broth without puromycin. Open symbols indicate when cells were sampled for assays of resting cells shown in Fig. 6. Error bars indicate standard deviations for 3 independent measurements. In many cases, the error bars were smaller than the symbols and are not shown.

**Fig. 4.**
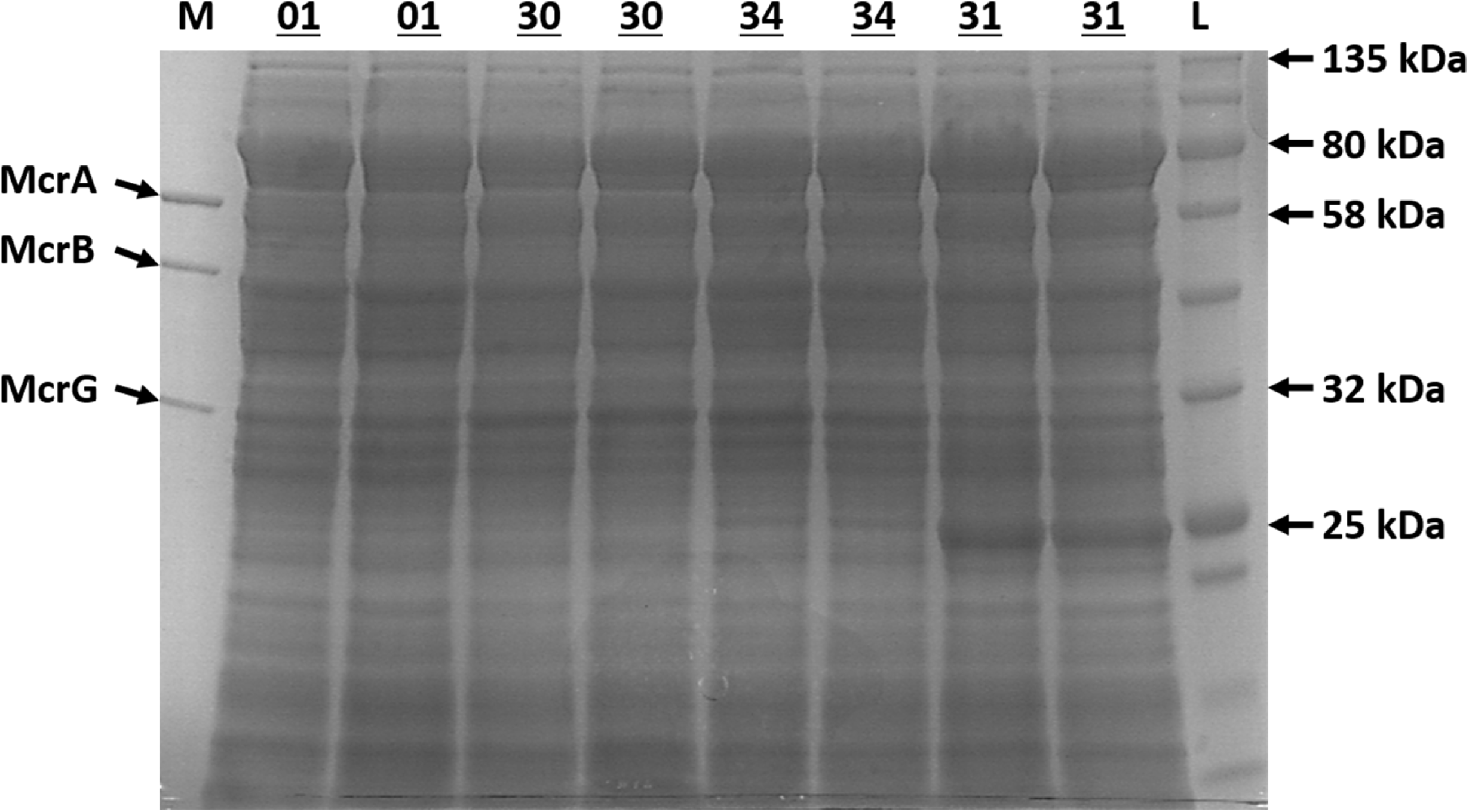
Mcr expression in the *∆mmp10* mutant and complementation strains. SDS PAGE profiles for strains: 01, S0001 (the wild-type); 30, S0030 (the deletion mutant); 34, S0034 (the *PhmvA-∆mmp10* complement); and 31, S0031 (the *Pnat-∆mmp10* complement). M, a purified wild-type his-tagged Mcr. L, protein ladder (NEB #P7712). Although only duplicate samples are shown here due to space, a third sample was also run on a different SDS gel that showed the identical pattern. Quantification of the Mmp10 was not possible because Mmp10 (48.9 kDa) and McrB (46.6 kDa) have nearly the same size and were not separated.

However, one band at the position of ~25 kDa showed substantial increase in intensity in strain S0031 (*p* < 0.0001 by One-way ANOVA). The relative abundances were 2.0 ± 0.1 % (average and standard deviation of three measurements, S0001), 1.6 ± 0.5 % (S0030), 2.8 ± 0.2 % (S0034) and 7.8 ± 0.7 % (S0031), respectively. MALDI sequencing analysis of the band from S0031 revealed that it was the puromycin N-acetyltransferase, which was encoded on the complementation plasmid and provided puromycin resistance.

The relative transcription rates of the *Pnat* and *PhmvA* promoters were also examined using an mCherry reporter system in strains S0032 and S0033, respectively. The mCherry fluorescence for S0032 was around 30-fold lower than S0033, i.e., arbitrary fluorescence values OD^−1^ mL^−1^ were 19.3 ± 1.5 (average and standard deviation of three measurements) versus 618 ± 40, respectively. Therefore, the differences in growth and methanogenesis observed between strains S0031 and S0034 could at least in part be attributed to the differences in expressional levels of *mmp10* between the two strains.

Collectively, these results suggested that the poor growth of the complementation strains was due to pleiotropic effects unrelated to the *mmp10* deletion. To test this hypothesis, the complementation plasmids were transformed into the wild type strain S0001, resulting in strains S0035 and S0036. Both strains grew poorly in comparison to the wild type and *∆mmp10* deletion strains (**Fig. 5A**). Moreover, the inhibition was even more profound than the corresponding complementation strains S0031 and S0034 in the *∆mmp10* background (**Fig. 5B**). Thus, the poor growth of the complementation strains appeared to be unrelated to their effect of Mcr methylation.

**Fig. 5.**
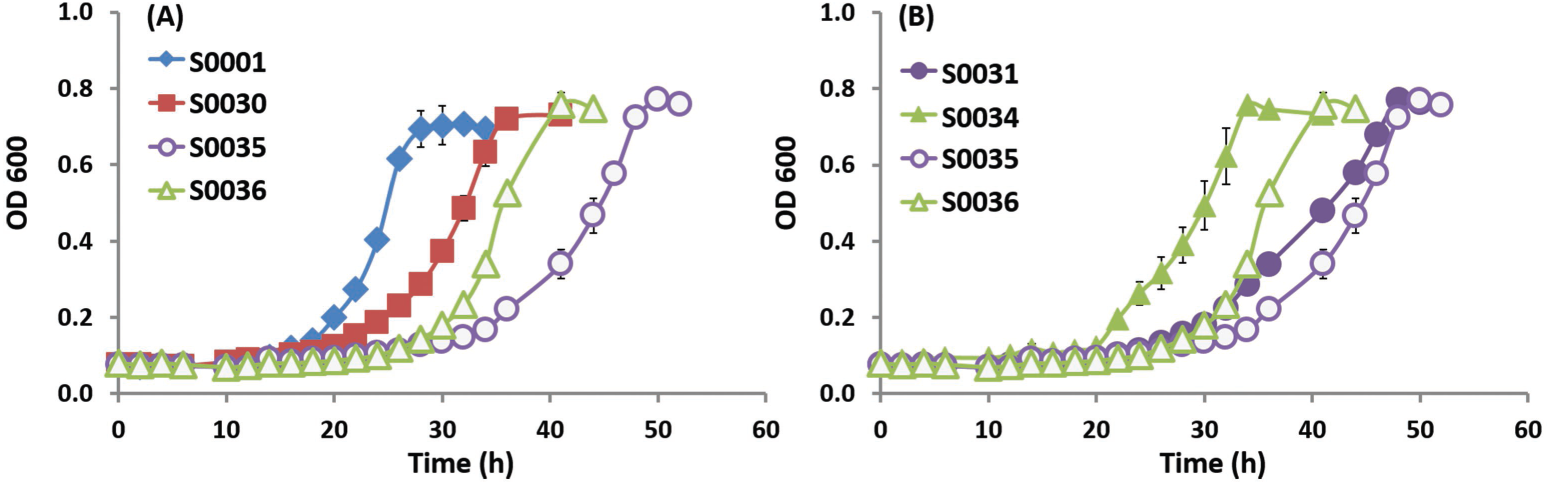
Growth of *M. maripaludis* strains with the complementation plasmids in a wild-type background. **(A)** Growth of both S0035 (S0001 + *Pnat-mmp10*) and S0036 (S0001 + *PhmvA-mmp10*) were inhibited compared to S0001 (the wild-type) and S0030 (the deletion mutant). **(B)** Growth inhibition for both S0035 and S0036 were more severe than the corresponding complementation strains with the plasmids in the *Δmmp10* background; S0031 (S0030 + *Pnat-mmp10*) and S0034 (S0030 + *PhmvA-mmp10*). Cultures were grown in 5 mL of complex formate broth without puromycin in 28 mL Balch tubes. Error bars indicate standard deviations for 3 independent measurements. In many cases, the error bars were smaller than the symbols and are not shown.

### Mcr activity in vivo

The Mcr from *M. maripaludis* is very unstable even in cell extracts, and greater than 90 % of the activity of resting cells is lost within one hour even under the conditions that stabilize the enzyme from *Methanothermobacter marburgensis* (unpublished observations). Therefore, the rates of methanogenesis of the *∆mmp10* mutant were compared to that of the wild-type in whole cells. Because the rates of methanogenesis by whole cells vary with the culture phase (31), the accumulation of methane in the culture headspace was followed throughout growth, and the rate of methanogenesis was calculated from the increases in methane in the culture headspace (**Fig. 6A**). By this measure, the rates of methanogenesis by the *∆mmp10* strain S0030 never exceeded 60% of the wild type rate. To confirm that these results were not due to fluctuations in the medium composition during growth, the rates of methanogenesis were also measured in resting cells after washing and resuspension in fresh medium (**Fig. 6B**), where a 60% reduction in the maximal rates of methanogenesis was observed in the *∆mmp10* strain S0030. These results confirmed that the rate of methanogenesis was severely impaired in the *∆mmp10* mutant.

**Fig. 6.**
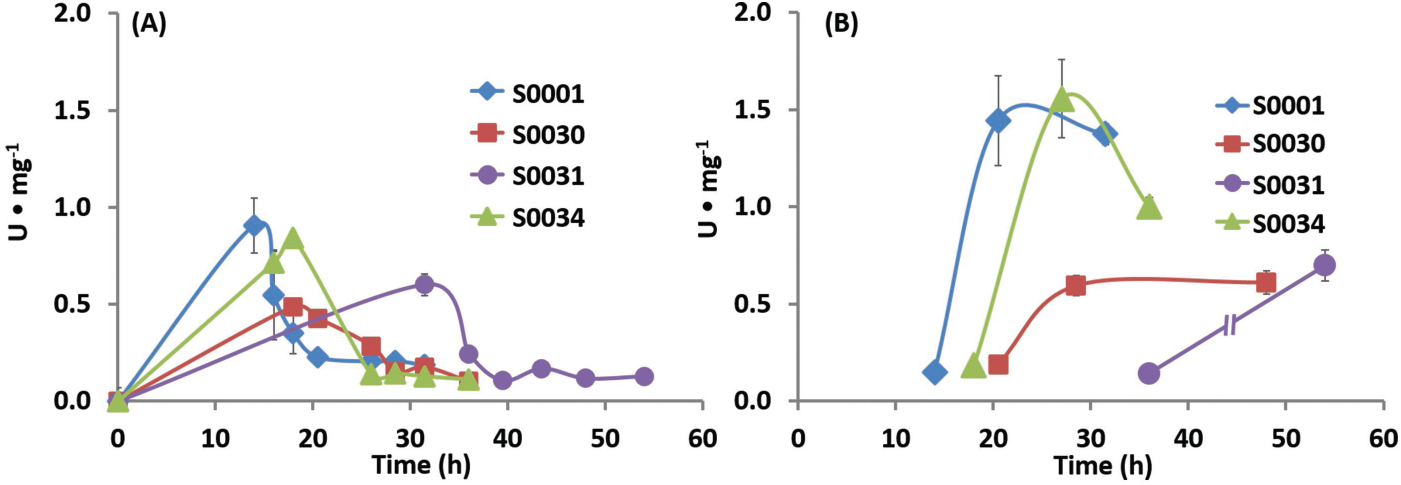
Cellular rate of methanogenesis in the *∆mmp10* mutant and complementation strains. **(A)** Rates of CH_4_ production during growth for the cultures shown in Fig. 3. The rates were calculated from the increases in CH_4_ in the headspace of the cultures. The maximal rates were 0.91 ± 0.14 (S0001), 0.49 ± 0.03 (S0030), 0.60 ± 0.07 (S0031), and 0.84 ± 0.06 (S0034) U•mg^−1^. **(B)** Specific CH_4_ production rates for resting cells sampled from cultures shown in the open symbols in Fig. 3. The maximal rates were 1.44 ± 0.23 (S0001), 0.61 ± 0.06 (S0030), 0.70 ± 0.08 (S0031), and 1.55 ± 0.20 (S0034) U•mg^−1^. Error bars indicate standard deviations for 3 independent measurements. U•mg^−1^ indicates μmol CH_4_ min^−1^ mg^−1^ dry weight.

In the complementation strains, the rate of methanogenesis was restored to the wild-type level in strain S0034, which possessed *mmp10* under the control of the strong *PhmvA* promoter, but the rate of methanogenesis for S0031, which possessed the *Pnat* promoter, was similar to that of the deletion mutant (**Fig. 6A and 6B**). In a separate experiment, the rates of methanogenesis for strains S0035 and S0036, which possessed the complementation plasmids in the wild-type background, were comparable to the wild-type level (data not shown).

## Discussion

In this study, bioinformatic analyses identified the *M. maripaludis* MMP1554 gene encoding Mmp10 as a candidate methyltransferase for PTMs of the Mcr. To test this hypothesis, a *∆mmp10* mutant was constructed, and the methylation of Arg^275^ in the active site region of Mcr was found to be lost. This PTM was restored when the mutant was complemented with *mmp10* expressed from plasmids, further supporting the bioinformatics prediction. However, even though Mmp10 contains structural features expected for a methyltransferase, these results do not eliminate the possibility that other proteins are required, and biochemical studies of Mmp10 will be required to confirm its activity. Because all other characterized arginine methylations take place at the guanidino nitrogen atoms instead of the C-5 (32), more information about the mechanism and role of Mmp10 is of great interest.

In resting cells, the rate of methanogenesis was reduced by 40-60 % in the *∆mmp10* mutant, suggesting a role of the Arg methylation in modulating Mcr activity. These results are consistent with previous Tn-seq experiments which demonstrated that, although *mmp10* was not an essential gene, transposons insertions led to a decrease in fitness (33). Thus, it would not be surprising if the unmethylated Mcr retained only partial activity.

Two contrasting models can be envisioned for the loss in activity in the absence of the Arg PTM. First, the Arg methylation could be an important structural feature of the active site and play an important role in catalysis. Presumably, loss of this methylation may disorientate its neighboring R^274^ at the active site and disrupt the interactions between the R^274^ and coenzyme B, which could have negative catalytic consequences (Fig. 2). Second, it could also play a role in Mcr assembly. In most methanogens, *mmp10* and *mcr* are divergently transcribed, suggesting that their expression is coordinated. Thus, during translation, protein folding may be coordinated with the PTMs, which may occur co-translationally as the nascent chain emerges from the ribosome (34, 35). This model is consistent with the observations that all the PTMs are deeply embedded within the native enzyme and likely to occur before insertion of coenzyme F_430_ and the complete folding of the enzyme (1, 9). Thus, the low rates of methanogenesis of the *mmp10* deletion mutant could be a consequence of Mcr misfolding rather than a direct effect on catalysis. Currently, it is not possible to distinguish between these possibilities in the absence of a reliable *in vitro* assay. Moreover, they are not necessarily mutually exclusive, and the Arg PTM may play multiple roles in Mcr activity.

In strains S0031 or S0034, complementation of the deletion with Mmp10 expressed on a plasmid with either its native promoter (*Pnat*) or a strong constitutive promoter (*PhmvA*) failed to restore wild-type growth even though the Mcr was fully methylated. This growth inhibition was reproduced in strains S0035 and S0036, where the same expression plasmids were instead transformed into the wild-type background. Therefore, both expression plasmids were inhibitive of growth. Thus, these results suggest that the failure of complementation to fully restore the growth phenotype could have resulted from effects independent of the effect on Mcr activity. Both methanogenesis and growth were substantially reduced in strain S0031, where the mutation was complemented with *mmp10* under control of the weak *Pnat* promoter. *mmp10* and *mcr* are divergently transcribed, and the *Pnat* and *Pmcr* promoter regions overlap. Thus, it was possible that cloning *Pnat* might have inhibited Mcr expression. However, the levels of Mcr found in whole cells were similar in this complement, the wild type and other strains. Thus, the lower methanogenesis activity was not a result of a direct effect on Mcr expression. Alternatively, *Pmcr* is also used to drive expression of puromycin N-transacetylase in the *pac* cassette of the complementation plasmids, and strain S0031 contained three-fold higher levels of this protein than S0034, which contained *PhmvA*. Thus, the poor growth and methanogenesis of strain S0031 could have been a consequence of the high expression of puromycin N-acetyltransferase.

Although the rate of methanogenesis in the *PhmvA* complementation strain S0034 was similar to that of wild type, the growth was somewhat slower. Because Mmp10 is probably highly expressed in this strain, it is possible that other proteins might have been post-translationally modified. SAM-dependent protein methyltransferases are known to specifically recognize the amino acid sequences flanking the amino acid to be methylated (9, 36, 37). Thus, the *M. maripaludis* proteome was searched *in silico* for the potential methylation consensus sequence PxRR^275^(A/S)R(G/A). Although an identical match was not found, one partial hit of RRSRG was identified in MMP0140, which encodes a putative hydrogenase maturation factor. Importantly, hydrogenase is a key enzyme in methanogenesis, and this gene is likely essential for growth (33, 38, 39). Therefore, spurious methylation of MMP0140 might well inhibit growth, and further investigation will be needed to address this possibility.

In conclusion, this study identified a gene required for the Arg PTM of Mcr, and a similar strategy that combines bioinformatic and experimental approaches may be employed in future gene discovery for the remaining methanococcal PTMs. The reduction of methanogenesis of the *mmp10* deletion mutant suggest that this PTM influences catalysis. However, the complex phenotype of the complementation strains suggests that *mmp10* may play additional roles beyond affecting Mcr activity, either in Mcr assembly or the PTM of other genes. The absence of *mmp10* in certain methanogens, such as *Methanomassiliicoccales* and the *Candidatus* ‘Bathyarchaeota’ and ‘Verstraetearchaeota’ phyla, also suggests that the PTMs of Mcr are more diverse than previously anticipated.

## Materials and Methods

### Strains and culture conditions

Strains used in this study are listed in **Table 3**. The complex formate broth and solid medium for cultivation and transformation of *M. maripaludis* have been described previously, except that cysteine hydrochloride was replaced with equal amount of coenzyme M and the Fe(NH_4_)_2_(SO_4_)_2_ stock was replaced by a more diluted FeSO_4_ stock to reduce the final iron concentration from 35.1 μM to 10.9 μM (11, 31, 33, 40). This avoided small amounts of precipitates that formed occasionally, possibly due to the formation of iron sulfide. Puromycin (2.5 μg mL^−1^) and 8-azahypoxanthine (0.25 mg mL^−1^) were added from anaerobic, sterile stocks as needed. All *M. maripaludis* cultures were grown at 37 °C with 15 Psi of N2/CO2 in the headspace. Growth curves were followed by measuring optical densities of 600 nm. The cell dry weight was calculated from the optical densities based on previously published calibration curves (31, 41).

**Table 3.**
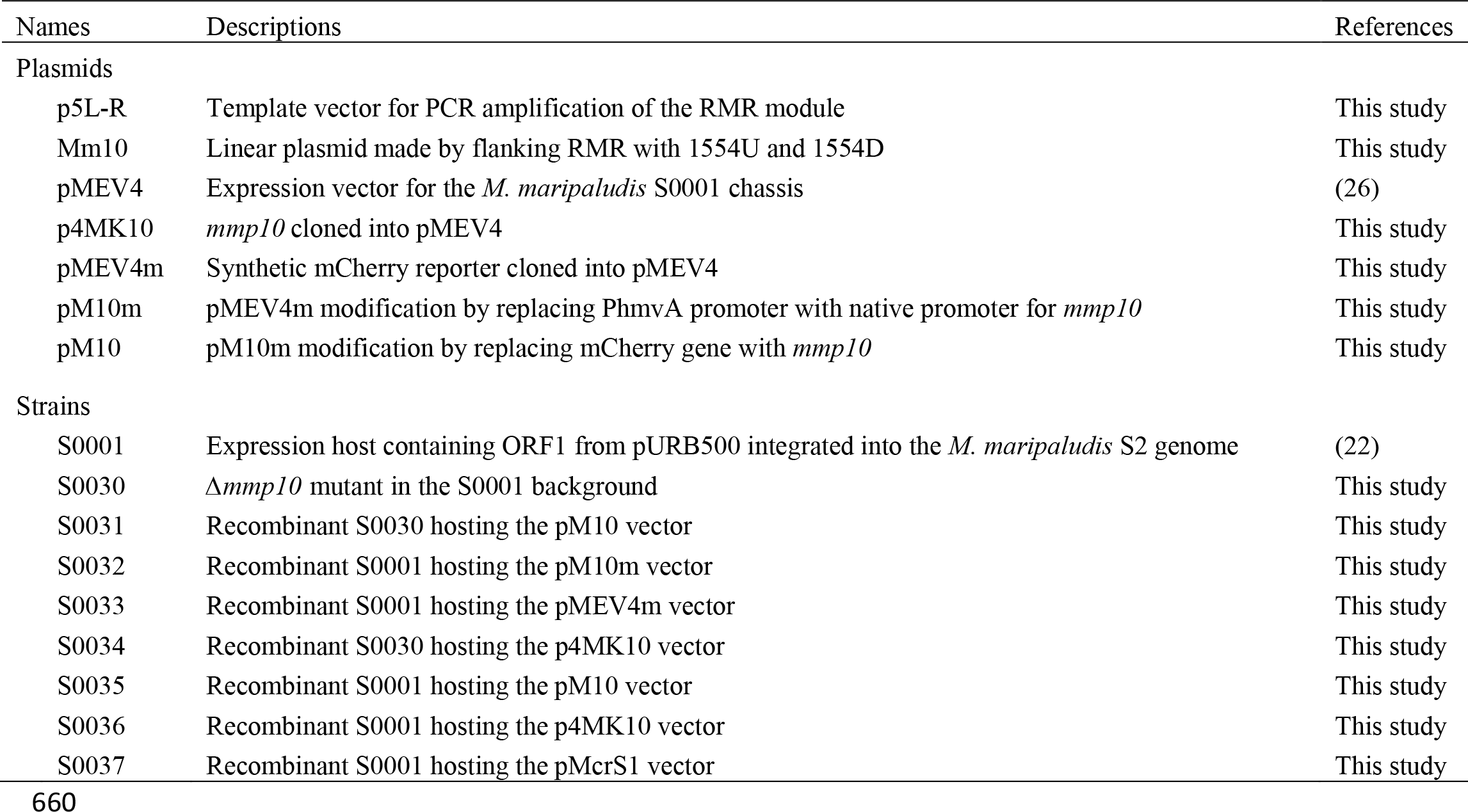
Plasmids and Microbial Strains

| Names | Descriptions | References |
| --- | --- | --- |
| Plasmids |  |  |
| p5L-R | Template vector for PCR amplification of the RMR module | This study |
| Mm10 | Linear plasmid made by flanking RMR with 1554U and 1554D | This study |
| pMEV4 | Expression vector for the *M. maripaludis* S0001 chassis | (26) |
| p4MK10 | *mmp10* cloned into pMEV4 | This study |
| pMEV4m | Synthetic mCherry reporter cloned into pMEV4 | This study |
| pM10m | pMEV4m modification by replacing PhmvA promoter with native promoter for *mmp10* | This study |
| pM10 | pM10m modification by replacing mCherry gene with *mmp10* | This study |
| Strains |  |  |
| S0001 | Expression host containing ORF1 from pURB500 integrated into the *M. maripaludis* S2 genome | (22) |
| S0030 | $\Delta mmp10$ mutant in the S0001 background | This study |
| S0031 | Recombinant S0030 hosting the pM10 vector | This study |
| S0032 | Recombinant S0001 hosting the pM10m vector | This study |
| S0033 | Recombinant S0001 hosting the pMEV4m vector | This study |
| S0034 | Recombinant S0030 hosting the p4MK10 vector | This study |
| S0035 | Recombinant S0001 hosting the pM10 vector | This study |
| S0036 | Recombinant S0001 hosting the p4MK10 vector | This study |
| S0037 | Recombinant S0001 hosting the pMcrS1 vector | This study |

### Plasmids and strains construction

Plasmids and PCR primers are listed in **Table 3** and **Table S3**. Cloning was performed in *E. coli* Top10, and selection was conducted with 100 μg mL^−1^ ampicillin. Genomic DNA extraction, DNA purification, and plasmid isolation were done with Zymo Research kits D6005, D4001/4003, and D4020, respectively, according to the manufacturer’s instructions. All PCR reactions were performed with the Phusion^®^ High-Fidelity DNA Polymerase (NEB, M0530S).

A linear DNA fragment Mm10 was constructed for performing markerless deletion of *mmp10* from the wild-type strain S0001, producing strain S0030. This strategy is described in detail in Supplementary Methods. Upon transformation, genomic integration of the Mm10 construct through homologous recombination was positively selected with puromycin. Colonies of the selected transformants were subjected to PCR amplification using primers 1554-F/R to confirm complete integration. Removal of the markers was achieved by selection with azahypoxanthine for a second homologous recombination between the two repetitive elements or RE. The identity of colonies of the *∆mmp10* mutant were confirmed by PCR amplification using primers 1554-F/R to confirm complete removal of the markers (**Fig S11A**). This conclusion was further confirmed by the sensitivity of the mutant to puromycin and the inability of primers 1554w-F/R, which target an internal region of *mmp10*, to yield a product. As a positive control, PCR amplification under the same conditions detected 0.1% of wild-type DNA when mixed with the mutant DNA (**Fig S11B**).

Expression plasmid vectors pM10 and p4MK10 were made to complement the *∆mmp10* in strain S0030, producing strains S0031 and S0034, respectively. The same two plasmids were also transformed into the wild-type strain S0001, resulting in strains S0035 and S0036, respectively. Expression of *mmp10* was under the control of the predicted native promoter *Pnat* in pM10. Expression of *mmp10* was under the control of the constitutive *PhmvA* promoter in p4MK10. The strength of *Pnat* and *PhmvA* promoters were quantified in strains S0032 and S0033, which harbored vectors pM10m and pMEV4m, respectively. The pMEV4m possessed the mCherry reporter under control of *PhmvA* and enabled gene expression to be quantified by following the mCherry fluorescence (Lyu and Whitman, unpublished observation). In pM10m, mCherry expression was controlled by *Pnat*. To make pM10m, two PCR products were made, digested with *Eco*RI and *Nde*I, and ligated by T4 DNA ligase (NEB, M0202S). The first PCR product was amplified from S0001 genomic DNA with primers 1554P-F/R, producing *Pnat*. The second PCR product was amplified from the pMEV4m plasmid with primers 4mp-F/R, producing the vector backbone with the *PhmvA* removed. Similarly, another two PCR products were made, digested with *Nde*I and *Pst*I, and ligated to make pM10. The first PCR product was amplified from S0001 genomic DNA with primers 1554c-Fa/R, producing the complete *mmp10* gene. The second PCR product was amplified from the pM10m plasmid DNA with primers p4Brk-F and 1554P-R, producing the native promoter and vector backbone with the mCherry gene removed. The p4MK10 was made by cloning *mmp10* into pMEV4 at the *Spe*I and *Pst*I sites. Specifically, *mmp10* was amplified from S0001 genomic DNA with primers 1554c-Fb/R, and the product was digested with *Xba*I and *Pst*I before ligation into the pMEV4 backbone. The pMEV4 backbone was amplified from the pMEV4 plasmid DNA with primers p4Brk-F/R.

Another expression vector pMcrS1 was constructed to express the wildtype *mcr* operon fused with a 6×-histag under control of the *PhmvA* promoter in strain S0001, resulting in a new strain S0037. Details regarding this construction will be reported elsewhere.

### Fluorescence quantification

Triplicate cultures were grown to an absorbance of 0.4~0.6 in the presence of puromycin. Cultures of 2 mL were harvested by centrifugation at 17, 000 g x 1 min, resuspended in 200 μL of PIPES-K buffer (25 mM and pH 6.8), and frozen at −80 °C overnight. For activating the mCherry, cell extracts were thawed and incubated overnight in air with shaking at 30 °C in the dark. The cell extracts were cleared by centrifugation at 17, 000 g x 1 min before measuring the fluorescence of the supernatant using Qubit 2.0 with excitation at 600-645 nm and emission at 665-725 nm.

### Purification of methyl-coenzyme M reductase

Cultures were grown in 1.5 L of complex formate broth to an absorbance of about 0.8. Cells were freshly harvested aerobically by centrifugation at 17,700 x g for 15 min at room temperature. Cells were resuspended in about 2 mL of Mcr buffer g^−1^ wet weight and stored at -20 ^°^C. The Mcr buffer contained 10 mM Ti(III) citrate, 10 mM coenzyme M, 0.1 mM methylviologen in 150 mM monosodium phosphate (pH 8.0) buffer. The cells in the Mcr buffer were thawed and remained on ice during sonication, which was conducted with a W-380 sonicator (Heat Systems-Ultrasonics, Inc) for 20 cycles of 5 s bursts with the output set at 5 and duty cycle set at 90 %. The lysate was then centrifuged at 17,000 x g for 5 min to remove cell debris. To achieve 50 % (NH_4_)2SO_4_ saturation, an equal volume of a saturated (NH_4_)2SO_4_ solution was added to the supernatant. After about 30 min, the precipitate was collected by centrifugation at 17,000 x g for 5 min and discarded. Additional (NH_4_)2SO_4_ powder was added to achieve 100 % saturation. After about 30 min, the precipitate was collected by centrifugation and resuspended in 4 mL of buffer A [1 mM coenzyme M in 10 mM Tris-HCl, pH 7.6]. The resulting proteins were desalted by concentration on an Amicon Ultra -4 centrifugal filter (Millipore, 10-kDa cutoff) by centrifugation at 7,500 x g for 15 min at 4 ^°^C and resuspension in the same buffer. The desalted and concentrated proteins were resuspended with 2.5 mL buffer A and loaded onto a Q-sepharose XK16 anion-exchange column equilibrated with the same buffer. The protein was eluted using an Akta fast protein liquid chromatograph (FPLC) system (GE healthcare) with a linear gradient of 0 % to 100 % buffer B [buffer A plus 1 M NaCl]. A similar protocol was used for purification of the his-tagged recombinant Mcr, except that coenzyme M was removed from the buffers and the ammonium sulfate precipitation was replaced by a Ni-Sepharose column (GE Healthcare) step, where protein was eluted using a linear imidazole gradient from 0 % to 100 % before the ion-exchange chromatography with Q-Sepharose. The Mcr subunits were separated by SDS PAGE and stained with AcquaStain (Bulldog Bio) for 2 to 10 min until protein bands appeared. The gel was then washed with ddH2O, and a slice of gel containing the McrA subunit was excised and destained twice with 30 % ethanol before being processed for mass spectrometry.

### Mass spectrometry analysis

For trypsin digestion, the gel bands were sliced into small pieces and rinsed twice with 50% acetonitrile/20 mM ammonium bicarbonate (pH 7.5~8.0). Proteins in the gel pieces were then alkylated by 50 mM iodoacetamide in 20 mM ammonium bicarbonate for 1 hour in the dark. The gel pieces were rinsed twice with 50 % acetonitrile in 20 mM ammonium bicarbonate, dehydrated by adding 100% of acetonitrile and dried by a SpeedVac. Then small amounts of trypsin solution (Promega, 0.01μg μL^−1^ in 20 mM ammonium bicarbonate) were added until the gel pieces totally absorbed the solution. The tubes were placed in an incubator at 37 °C overnight. The tryptic peptides were extracted twice from gel pieces with 50 % acetonitrile in 0.1 % formic acid. The extracts were then combined and taken to dryness on a SpeedVac. A similar protocol was used for pepsin digestion, except that the pH was adjusted to ~2 with 0.04 M HCl and the digestion was performed for 48 h.

The mass spectrometry analyses were performed on a Thermo-Fisher LTQ Orbitrap Elite Mass Spectrometer coupled with a Proxeon Easy NanoLC system (Waltham, MA) located at the Proteomics and Mass Spectrometry Facility, University of Georgia. The peptides were loaded onto a reversed-phase column (Dionex PepMap 100 C8, or 100 μm id column/emitter self-packed with 200 Å 5 μM Bruker MagicAQ C18 resin) and then eluted into the mass spectrometer. Briefly, the two-buffer gradient elution at a flow rate of 500 nL min^−1^ (0.1% formic acid as buffer A and 99.9 % acetonitrile with 0.1 % formic acid as buffer B) started with 5 % B for 2 min, then increased to 25 % B in 60 min, to 40 % B in 10 min, and finally to 95% B in 10 min.

The data-dependent acquisition (DDA) method was used to acquire MS data. A survey MS scan was acquired first, and then the top 5 ions in the MS scan were selected for CID and HCD MS/MS experiments. Whenever necessary, ETD was used instead of CID for better identification of post-translational modifications (42). MS and MS/MS scans were acquired by Orbitrap at the resolutions of 120,000 and 30,000, respectively. Data were acquired using Xcalibur software (version 2.2, Thermo Fisher Scientific). Protein identification and modification characterizations were performed using Thermo Proteome Discoverer (version 1.3/1.4) with Mascot (Matrix Science) or SEQUEST (Thermo) programs.

### Resting cell rates of methanogenesis

To follow CH_4_ production during growth, cultures were inoculated into 20 mL of formate medium without puromycin in 210 mL anaerobic bottles with fused-in side arms for convenient measurement of cell densities unless otherwise mentioned. The headspace was sampled for CH_4_ throughout growth. For assays of resting cells, 4 mL of culture were harvested at different growth stages by centrifugation at 2800 x *g* for 10 min inside an anaerobic chamber. Triplicate cell pellets were washed and resuspended in 1.6 mL of formate medium in a 4.6 mL anaerobic vial sealed with a butyl rubber stopper. Vials were flushed immediately with an atmosphere of N2/CO2 for 30 s, and the assay was initiated by incubating at 37 °C. Headspace gas was withdrawn from the vials at intervals of 3-6 min for 30-50 min, and methane was detected with an SRI 8610-C gas chromatograph as described previously (31). One unit was defined as 1 μmol of CH_4_ produced per min.

During the growth experiment, CH_4_ production rates in the culture were corrected for the removal of cells during sampling. Specific growth rates during exponential growth were analyzed by linear regression of the logarithm of the optical density with time. In parallel, cell extracts from the culture were separated in an SDS gel to examine protein expression profiles (43). The relative abundance for any proteins of interest in the SDS gel was also estimated by integrating peak area for that band and compared to the total peak areas for the entire lane using ImageJ (44). All samples were prepared in triplicate, and mean values of the triplicates ± standard deviation were presented.

### Bioinformatic analyses

Distribution of the methanogenesis marker 10 gene family TIGR03278 across Archaea was assessed using the Integrated Microbial Genomes & Microbiomes Expert Review or IMG/M ER platform (45, 46). Not hosted by the IMG/M ER, ANME-1 (FP565147) and Verstraetearchaeota (GCA_001717005, GCA_001717035, GCA_001717015, GCA_001717085 and MAGU00000000) genome assemblies from Genbank were searched by BLAST for the marker 10 homologs from ANME-2 and *M. maripaludis*. Terminators were predicted by ARNold (47). Protein structural analysis was conducted with either EMBOSS 6.5.7 (48) or Protein Workshop (49). Unless otherwise mentioned, all other analyses were done with Geneious versions 8 and 10 (50).

## Author Contributions

The experiments were conceived and designed by LZ, CWC, ECD and WBW. They were executed by LZ, CWC, SH and RP. The manuscript was written by LZ, CWC and WBW.

## Acknowledgment

We thank Aynom Tsegay Misghina and Bhavana Nagareddy for their assistance in developing the p5L-R tool, the University of Georgia iGEM team for their contribution in developing the methanoccoccal mCherry reporter system, Hannah Bullock for assistance using the FPLC, and the Whitman lab members, especially Suet Yee Chong and Nana Shao, for assistance with crystal structure analysis. In addition, partial support was provided by the Office of the Vice-President for Research at the University of Georgia, the Korea Institute of Ocean Science and Technology, National Science Foundation grant MCB-1410102, and a contract from the U.S. Department of Energy.

